# Plant detection and counting from high-resolution RGB images acquired from UAVs: comparison between deep-learning and handcrafted methods with application to maize, sugar beet, and sunflower

**DOI:** 10.1101/2021.04.27.441631

**Authors:** Etienne David, Gaëtan Daubige, François Joudelat, Philippe Burger, Alexis Comar, Benoit de Solan, Frédéric Baret

**Affiliations:** INRAe, UMR EMMAH, Avignon, France; Arvalis – Institut du végétal, Avignon, France; ITB – Institut Technique de la Betterave, Paris, France; INRAe, UE Grandes Cultures Auzeville (GCA), Toulouse, France; Hiphen, 22b Rue Charrue, Avignon, France

**Keywords:** Plant counting, Deep-learning, Robustness, unmanned aerial vehicle, phenotyping

## Abstract

Progresses in agronomy rely on accurate measurement of the experimentations conducted to improve the yield component. Measurement of the plant density is required for a number of applications since it drives part of the crop fate. The standard manual measurements in the field could be efficiently replaced by high-throughput techniques based on high-spatial resolution images taken from UAVs. This study compares several automated detection of individual plants in the images from which the plant density can be estimated. It is based on a large dataset of high resolution Red/Green/Blue (RGB) images acquired from Unmanned Aerial Vehicules (UAVs) during several years and experiments over maize, sugar beet and sunflower crops at early stages. A total of 16247 plants have been labelled interactively on the images. Performances of handcrafted method (HC) were compared to those of deep learning (DL). The HC method consists in segmenting the image into green and background pixels, identifying rows, then objects corresponding to plants thanks to knowledge of the sowing pattern as prior information. The DL method is based on the Faster Region with Convolutional Neural Network (Faster RCNN) model trained over 2/3 of the images selected to represent a good balance between plant development stage and sessions. One model is trained for each crop.

Results show that simple DL methods generally outperforms simple HC, particularly for maize and sunflower crops. A significant level of variability of plant detection performances is observed between the several experiments. This was explained by the variability of image acquisition conditions including illumination, plant development stage, background complexity and weed infestation. The image quality determines part of the performances for HC methods which makes the segmentation step more difficult. Performances of DL methods are limited mainly by the presence of weeds. A hybrid method (HY) was proposed to eliminate weeds between the rows using the rules developed for the HC method. HY improves slightly DL performances in the case of high weed infestation. When few images corresponding to the conditions of the testing dataset were complementing the training dataset for DL, a drastic increase of performances for all the crops is observed, with relative RMSE below 5% for the estimation of the plant density.

## 1 Introduction

Measuring accurately traits is essential for numerous applications in agronomy, such as breeding or new farm management strategies evaluation. Plant density at emergence is a main yield component particularly for plants with reduced tillering or branching capacities such as maize, sugar beet and sunflower. The plant density at emergence is controlled by the seeding density and the emergence rate. Further, the seeding pattern defined by the distance between row and between plants influences the competition between plants and possibly with weeds. In addition to the estimation of plant density, the position of each plant can be documented to describe the local competitive environment (Godwin and Miller, 2003). For agronomical or phenotyping experiments, the plant density is mainly used to evaluate the quality of each microplot with consequences on the whole trial. It is also used by farmers to decide to stop spending resources to grow the crop in case of too low density or too much heterogeneity. Plant density is considered as an agronomical trait in some widely used ontology (Shrestha et al., 2012), despite not being directly governed by the genotype, as it results from the seeding density, seed vigor and the emergence conditions.

Plant density is assessed manually in current breeding programs. Operators count plants in the field over a limited sampling area, usually less than 1 square meter, since this process is tedious, time-consuming, and therefore expensive. Consequently, this traditional method can lead to significant uncertainties due to the limited representativeness of the sampled area and possible human errors. Further, the position of plants is generally not documented because it would be even more tedious to measure each plant location.

The recent technological advances of plant phenotyping solutions including Unmanned Aerial Vehicles (UAV), sensors, computers, and image processing algorithms, offer potentials to develop alternative methods to the manual counting. Several authors already reported accurate estimates of plant or organ counting and density from RGB images (Table 1). Plants or organ can be characterized either with machine learning (ML) algorithms where standard local image features are extracted and a used in a supervised classification to identify the objects of interest (Guo *et al*., 2018; Fernandez-Gallego *et al*., 2019). Handcrafted (HC) methods rely on expert knowledge to compute the pertinent features in a process known as “feature engineering” and use them to identify the objects of interest. Most of them belong to the Object Based Image Analysis (Josue Nahun Leiva et al., 2017; Koh et al., 2019; Torres-Sánchez et al., 2015; Varela et al., 2018; Zhao et al., 2018). The identification process can be done based also on the expert knowledge (Gnädinger and Schmidhalter, 2017; Jacopin et al., 2021; T. Liu et al., 2016) or by calibrating a statistical model over a training dataset (Calvario et al., 2020). More recently, approaches based on deep-learning (DL) have been proposed. The features are automatically extracted from the image and then used to identify and localize the individual objects of interest ((Lin and Guo, 2020; Liu et al., 2020; Madec et al., 2019; Quan et al., 2019)). However, these features can also be used to estimate directly the density of objects through a regression (Ribera et al., 2017; Valente et al., 2020; Xiong et al., 2019). Localization, is more popular (78% of the studies in Table 1) in plant phenotyping as it documents the sowing heterogeneity including missing plants, allowing to explore the competition between plants as outlined earlier. DL based methods are being common now to detect plant and organ and represent almost 30% of the localization studies (Table 1). Madec et al. (Madec et al., 2019) demonstrated that the Faster RCNN DL model (Ren et al., 2015) provides accurate localization of wheat ears with higher robustness than previous methods, including direct regression method. A higher heritability than that of manual counting was also reported. More recently, (Lin and Guo, 2020; Liu et al., 2020) applied similar strategies to locate plant and organ from UAV images. DL applications to plant phenotyping are supervised learning methods, requiring large and diverse labelled datasets to converge to a generic solution. The recent progress in DL applied to detection/localization tasks beneficiated from the availability of large image collections such as ImageNet (Deng et al., 2009) and COCO Dataset (Lin et al., 2014) that are used to pre-train the DL model.

**Table 1:** **Comparison of the different approaches used for plant and organ counting referenced in the literature**. ^**1**^ **random selection of samples for training and testing;** ^**2**^**No proper calibration;** ^**3**^**Calibrated with synthetic data;** ^**4**^**Testing is made on two sessions, one session being already used for training**

| # | Study | UAV | Crop | Object | Sessions | Localiz<br>ation | Method | Test<br>independency |
| --- | --- | --- | --- | --- | --- | --- | --- | --- |
| 1 | (Guo et al., 2018) | Yes | Sorghum | Head | 1 | Yes | ML | No<sup>1</sup> |
| 2 | (Fernandez-Gallego et al., 2020) | yes | Wheat | Plant | 5 | yes | ML | No<sup>1</sup> |
| 3 | (T. Liu et al., 2016) | no | Wheat | Plant | several | yes | HC | Yes<sup>2</sup> |
| 4 | (Gnädinger and Schmidhalter, 2017) | yes | Maize | Plant | 1 | yes | HC | yes<sup>2</sup> |
| 5 | (Jacopin et al., 2021) | yes | Sunflower | Plant | 1 | yes | HC | Yes<sup>3</sup> |
| 6 | (Calvario et al., 2020) | yes | Agave | Plant | 3 | yes | HC | No |
| 7 | (Torres-Sánchez et al., 2015) | yes | Maize<br>Sunflower<br>wheat | Plant | 6 | no | HC (OBIA) | Yes<sup>2</sup> |
| 8 | (Josue Nahun Leiva et al., 2017) | yes | Thuja | Plant | 3 | yes | HC (OBIA) | Yes<sup>2</sup> |
| 9 | (Varela et al., 2018) | yes | Maize | Plant | 2 | yes | HC (OBIA) | No<sup>1</sup> |
| 10 | (Zhao et al., 2018) | yes | Rapeseed | Plant | 2 | yes | HC (OBIA) | No<sup>1</sup> |
| 11 | (Koh et al., 2019) | Yes | Safflower | Plant | 2 | Yes | HC (OBIA) | No<sup>4</sup> |
| 12 | (Madec et al., 2019) | No | Wheat | Head | 2 | yes | DL | Yes |
| 13 | (Quan et al., 2019) | No | Maize | Plant | 10 | yes | DL | No<sup>1</sup> |
| 14 | (Ribera et al., 2017) | Yes | Sorghum | Plant | 2 | no | DL | No<sup>1</sup> |
| 15 | (Xiong et al., 2019) | Yes | Wheat | Head | several | no | DL | Yes |
| 16 | (Valente et al., 2020) | Yes | Spinach | Plant | 1 | no | DL | No<sup>1</sup> |
| 17 | (Liu et al., 2020) | Yes | Maize | Head | 2 | yes | DL | No<sup>1</sup> |
| 18 | (Lin and Guo, 2020) | Yes | Sorghum | Head | 2 | yes | DL | No<sup>1</sup> |
|  | This study | Yes | Maize<br>Sugar beet<br>Sunflower | Plant | 27 | yes | HC / DL | Yes |

However, Geiros et al. (Geirhos et al., 2020) raised the overfitting risk and the resulting lack of robustness associated with most DL algorithms. They can reach excellent performances for datasets like those used for their calibration, while often failing when applied to cases different from the training dataset. In comparison, HC methods are based on expert knowledge which select the main features to identify the target objects. This reduces the risk of overfitting but can hardly account for all the specific cases. On the 11 methods listed (Table 1) that require a training dataset, only 3 (Koh et al., 2019; Madec et al., 2019; Xiong et al., 2019) proposed a proper evaluation framework where the training and the test datasets do not come from the same acquisition sessions. This questions the accuracy, scalability and robustness of HC and DL methods that was investigated in the case of liver disease (Lin et al., 2020), but not for the plant detection problem within phenotyping applications.

The objective of this study is to compare a HC approach based on the knowledge of the sowing and plant patterns and a DL approach based on object detection to localize plants and count them. This study includes three species (maize, sugar beet and sunflower) observed with a RGB (Red Green Blue) camera aboard a UAV during 27 acquisition sessions with plants at different development stages few weeks after emergence. This study appears therefore to be the most comprehensive one on the subject (Table 1), while keeping always the training and test datasets as independent as possible. Further, we will also propose to combine the DL approach with expert knowledge from the HC one.

## 2 Materials and methods

### 2.1 Dataset

#### 2.1.1 Experiments

The dataset used was acquired over maize, sugar beet and sunflower experiments from 2016 to 2019 in several experimental sites in France (Table 2). The sites cover a large diversity of agronomic conditions while managed with conventional tillage practices. However, some crop residues from the previous season can be observed on few microplots. Generally, few weeds were present in the microplots, except for some of them (Table 3). The sites include clay, brunisolic and limestone soil types (Table 2) with a variety of surface roughness and moisture. The soil color varies from gray to brown due to soil type, surface aspect and illumination conditions. Each site included an ensemble of microplots corresponding to many genotypes from which 3 to 12 were selected to get approximately 600 plants (Table 3). Some sites were flown several times (Table 2), corresponding to several acquisition sessions. This allows to get a larger variation in the crop development stage during image acquisition (Table 3). For maize, a total of 51 microplots was available from 9 acquisition sessions (Table 3) with contrasted microplot size, row spacing (0.3-1.1m), and plant density (5.1-11.2 plt.m^-2^). For sugar beet, a total number of 60 microplots was available from 9 acquisition sessions with microplot size, row spacing and plant density varying within a small range (Table 2). For sunflower, a total of 78 microplots was available from 9 acquisition sessions with a large variability of microplot size, row spacing, and plant density.

**Table 2:** Characteristics of the crops for the several sites considered.

| Crop | Site Name | Lat (°) | Long (°) | Year | Nb. sessions | Nb. microplots | Microplot width (m) | Microplot length (m) | Row spacing (m) | Plant density (plt.m<sup>-2</sup>) | Soil type |
| --- | --- | --- | --- | --- | --- | --- | --- | --- | --- | --- | --- |
| Maize | Menainville | 47.9 | 1.4 | 2016 | 1 | 6 | 2.2 | 7.0 | 1.10 | 5.1 | Clay |
|  | Nerac | 44.1 | 0.3 | 2016 | 1 | 8 | 1.6 | 7.0 | 0.80 | 8.5 | Clay |
|  | Villedieu | 47.8 | 1.5 | 2016 | 1 | 6 | 0.9 | 11.0 | 0.30 | 19.9 | Clay |
|  | Thenay | 47.3 | 1.2 | 2017 | 1 | 6 | 4.4 | 6.0 | 0.63 | 7.3 | Clay / Flint |
|  | Blois | 47.7 | 1.2 | 2019 | 1 | 7 | 1.7 | 7.0 | 0.83 | 9.5 | Brunisolic |
|  | Castetis | 43.4 | -0.7 | 2019 | 1 | 5 | 2.8 | 4.0 | 0.70 | 11.2 | Brunisolic |
|  | Ermine | 46.5 | -1.0 | 2019 | 1 | 4 | 3.2 | 5.5 | 0.80 | 8.6 | Limestone |
|  | Selommes | 47.7 | 1.2 | 2019 | 1 | 7 | 1.8 | 5.3 | 0.88 | 9.5 | Brunisolic |
|  | Pleinefougeres | 48.5 | -1.5 | 2020 | 1 | 2 | 3.2 | 11.0 | 0.80 | 7.7 | Brunisolic |
| Sugar beet | Bucy | 49.6 | 3.9 | 2017 | 2 | 7 | 1.4 | 6.2 | 0.45 | 11.1 | Loam |
|  | Charmont | 48.3 | 4.1 | 2017 | 1 | 7 | 1.4 | 5.5 | 0.45 | 11.1 | Limestone |
|  | Etienne | 49.2 | 4.3 | 2017 | 1 | 6 | 1.2 | 7.6 | 0.40 | 15.6 | Limestone |
|  | Memmie | 48.9 | 4.3 | 2017 | 2 | 6 | 1.4 | 7.6 | 0.48 | 10.8 | Limestone |
|  | Charmont | 48.3 | 4.1 | 2018 | 2 | 8* | 1.4 | 5.5 | 0.45 | 11.4 | Limestone |
|  | Memmie | 48.9 | 4.3 | 2018 | 1 | 6 | 1.4 | 7.6 | 0.45 | 11.4 | Limestone |
| Sunflower | Rivière | 43.5 | 1.5 | 2017 | 1 | 8 | 3.0 | 4.1 | 0.50 | 7.1 | Clay |
|  | Auzeville | 43.5 | 1.5 | 2018 | 2 | 3 | 3.3 | 9.5 | 0.55 | 6.1 | Clay |
|  | Auzeville | 43.5 | 1.5 | 2019 | 5 | 12 | 2.9 | 9.0 | 0.96 | 3.7 | Clay |
|  | Epoisses | 47.2 | 5.1 | 2019 | 1 | 4 | 2.4 | 10.0 | 0.60 | 5.1 | Limestone |

**Table 3:**
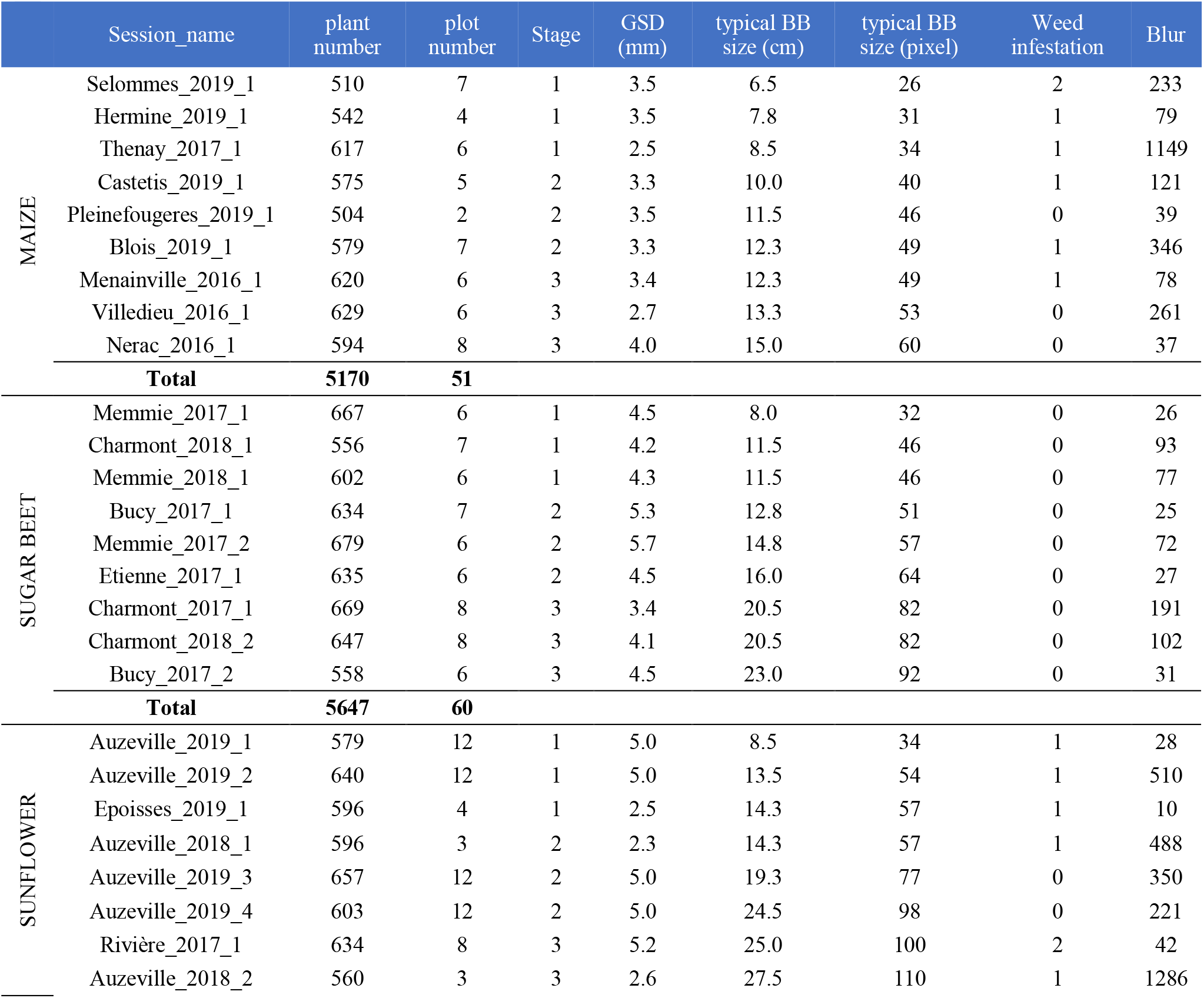
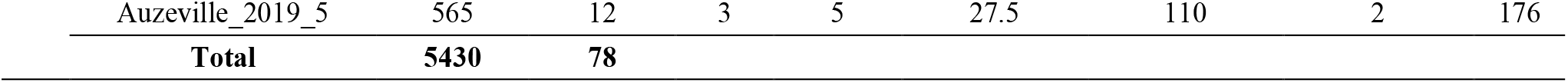
**Characteristics of each measurement sessions. For sugar beet, microplots from one session to another are the same. For sunflower the microplots considered change between sessions. The typical size of the BB for one session is computed as the square root of the mean area of all the BBs. The typical bounding box (BB) size in pixels is computed after up sampling the images at 2.5 mm resolution. The plant stage at the time of the session is quantified as: 1: early, 2: intermediate, 3: late. The correspondence with BBCH scale is provided as a table in the supplementary material section. The weed infestation is scored from 0 (no weed), from 0 (no weeds), 1 (less than 5% coverage), 2 (more than 5% coverage). The image blur is quantified by the average variance of the Laplacian: high blur results in low value of the variance of the Laplacian**.

|  | Session_name | plant number | plot number | Stage | GSD (mm) | typical BB size (cm) | typical BB size (pixel) | Weed infestation | Blur |
| --- | --- | --- | --- | --- | --- | --- | --- | --- | --- |
| MAIZE | Selommès_2019_1 | 510 | 7 | 1 | 3.5 | 6.5 | 26 | 2 | 233 |
|  | Hermine_2019_1 | 542 | 4 | 1 | 3.5 | 7.8 | 31 | 1 | 79 |
|  | Thenay_2017_1 | 617 | 6 | 1 | 2.5 | 8.5 | 34 | 1 | 1149 |
|  | Castetis_2019_1 | 575 | 5 | 2 | 3.3 | 10.0 | 40 | 1 | 121 |
|  | Pleinefougeres_2019_1 | 504 | 2 | 2 | 3.5 | 11.5 | 46 | 0 | 39 |
|  | Blois_2019_1 | 579 | 7 | 2 | 3.3 | 12.3 | 49 | 1 | 346 |
|  | Menainville_2016_1 | 620 | 6 | 3 | 3.4 | 12.3 | 49 | 1 | 78 |
|  | Villedieu_2016_1 | 629 | 6 | 3 | 2.7 | 13.3 | 53 | 0 | 261 |
|  | Nerac_2016_1 | 594 | 8 | 3 | 4.0 | 15.0 | 60 | 0 | 37 |
|  | **Total** | **5170** | **51** |  |  |  |  |  |  |
| SUGAR BEET | Memmie_2017_1 | 667 | 6 | 1 | 4.5 | 8.0 | 32 | 0 | 26 |
|  | Charmont_2018_1 | 556 | 7 | 1 | 4.2 | 11.5 | 46 | 0 | 93 |
|  | Memmie_2018_1 | 602 | 6 | 1 | 4.3 | 11.5 | 46 | 0 | 77 |
|  | Bucy_2017_1 | 634 | 7 | 2 | 5.3 | 12.8 | 51 | 0 | 25 |
|  | Memmie_2017_2 | 679 | 6 | 2 | 5.7 | 14.8 | 57 | 0 | 72 |
|  | Etienne_2017_1 | 635 | 6 | 2 | 4.5 | 16.0 | 64 | 0 | 27 |
|  | Charmont_2017_1 | 669 | 8 | 3 | 3.4 | 20.5 | 82 | 0 | 191 |
|  | Charmont_2018_2 | 647 | 8 | 3 | 4.1 | 20.5 | 82 | 0 | 102 |
|  | Bucy_2017_2 | 558 | 6 | 3 | 4.5 | 23.0 | 92 | 0 | 31 |
|  | **Total** | **5647** | **60** |  |  |  |  |  |  |
| SUNFLOWER | Auzeville_2019_1 | 579 | 12 | 1 | 5.0 | 8.5 | 34 | 1 | 28 |
|  | Auzeville_2019_2 | 640 | 12 | 1 | 5.0 | 13.5 | 54 | 1 | 510 |
|  | Epoisses_2019_1 | 596 | 4 | 1 | 2.5 | 14.3 | 57 | 1 | 10 |
|  | Auzeville_2018_1 | 596 | 3 | 2 | 2.3 | 14.3 | 57 | 1 | 488 |
|  | Auzeville_2019_3 | 657 | 12 | 2 | 5.0 | 19.3 | 77 | 0 | 350 |
|  | Auzeville_2019_4 | 603 | 12 | 2 | 5.0 | 24.5 | 98 | 0 | 221 |
|  | Rivière_2017_1 | 634 | 8 | 3 | 5.2 | 25.0 | 100 | 2 | 42 |
|  | Auzeville_2018_2 | 560 | 3 | 3 | 2.6 | 27.5 | 110 | 1 | 1286 |

|  |  |  |  |  |  |  |  |  |
| --- | --- | --- | --- | --- | --- | --- | --- | --- |
| Auzeville_2019_5 | 565 | 12 | 3 | 5 | 27.5 | 110 | 2 | 176 |
| **Total** | **5430** | **78** |  |  |  |  |  |  |

#### 2.1.2 Acquisition and labelling details

Image acquisition was carried out by UAVs embarking three different RGB cameras including the Sony Alpha 5100, Sony Alpha 6000, both with a resolution of 6024×4024 pixel, and the Zenmuse X7 (DJI) in the case of Epoisses site in 2019 with a resolution of 6016 × 4008 pixels. The cameras were fixed on a two axes gimbal to maintain the nadir view direction during the flight. The camera was set to speed priority of 1/1250 s to limit motion blur. The aperture and ISO were automatically adjusted by the camera. The camera was triggered by an intervalometer set at 1Hz frequency corresponding to the maximum value allowed to record the RGB images in JPG format on the memory card of the camera. Flight altitude above ground varied between 20 to 50m to get a ground sampling distance (GSD) between 2 mm and 5 mm per pixel (Table 3). The flight trajectory was designed to ensure more than 70% overlap between images across and along tracks. Ground control points were placed in the field and their coordinates were measured with a real-time kinetic GPS device ensuring an absolute centimetric accuracy of their position.

Agisoft Photoscan Professional software (Pasumansky, 2016) was used to align the images. The high overlap between the images and structure from motion algorithm permits to compute the position and orientation of the cameras. The pipeline described in Jin et al. (Jin et al., 2017) was then run to extract from each image the portion corresponding to the contained microplots, by extracting microplot thanks to a georeferenced plot map. Using the original images avoids the possible distortions and artefacts observed in the orthomosaic. Several extracts may represent the same microplot viewed from different positions of the UAV (Duan et al., 2016). For each microplot, the sharpest extract that contained the whole microplot is selected. For each session, a few microplots were selected for labelling (Table 2). Approximately 600 plants per session were labelled to ensure consistency across sessions which resulted in a total of 16247 labelled plants. Images were rescaled to match the best available GSD (2.5 mm, Table 3). This was necessary to control the apparent size of object, which can make the Deep Learning methods fail. Then all images were labelled using the coco-annotator tool (Brooks, 2019), an open source platform which allow the collaborative drawing of bounding box (BB) around each plant, which will be used as label. Six different operators contributed to the labelling. The labelling from one operator was always reviewed at least once by a different operator. The typical size of the BB for one session (Table 3) was computed as the square root of the mean area of all the BBs.

The plant development stage during the acquisition sessions was scored into three relative levels, where stages 1, 2 and 3 correspond respectively to early (few days after emergence), intermediate, and late stages (leaves start to fill the gap between plants). The correspondence between the stages for each crop, and their BBCH scale is presented in Table S1. The level of weed infestation (Table 3) was also visually evaluated from 0 (no weeds), 1 (sparse presence of weeds), 2 (infestation). The level of blurriness for each session (Table 3) was evaluated by calculating the average variance of the discrete Laplacian (Bansal et al., 2016), which is implemented in python with OpenCV.

### 2.2 Plant detection methods

#### 2.2.1 Handcrafted method

The method developed here is based on several assumptions: (1) the plants are green and can be accurately separated from the background; (2) plants are sown in rows relatively evenly spaced and parallel; (3) the weeds are mainly located in between the rows and are not too dominant; (4) plants are relatively evenly spaced on the row and are not too variable in shape and size. The method first extracts each single row and then identifies each individual plant on the row. All the parameters of our HC method are expressed in relative value to the row or plant spacing, to allow adaptation to a larger number of sowing patterns. This makes our method scalable to all our experimental conditions across the three species (Table 2 and table 3). The values of the parameters were set based on reasonable assumptions and were not calibrated on a dataset.

##### 2.2.1.1 Row extraction

The original RGB images are first transformed into a black and white one (BW) using the excess green index (Equation 1). Pixels are then assigned to the green (1) or background (0) classes using the ExG threshold value defined with the Otsu algorithm for each session (Otsu, 1975). Otsu algorithm is a method to perform automatic image thresholding based on the maximization of the class inter-variance. We used the implementation of python OpenCV library.

**Equation 1:** 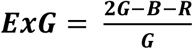. **R, G, B correspond respectively to the red, green and blue colours of the original image (Meyer and Neto, 2008)**

The Hough transform (Hough, 1962) is used to identify the main alignments corresponding to the rows and find their orientation. For each pixel assigned to green (1), several lines are drawn with different directions and for each line, the number of pixels it crosses is accumulated, allowing to find the orientation of the longest lines. We used Hough Transform implementation of python OpenCV library. The image is then rotated to display the rows horizontally (Figure 2). The number of green pixels in each line is computed to obtain a profile of green pixels across the rows. The peaks of the green pixel profiles are localized using the prior knowledge on row spacing (*Row_spacing_prior*) to prevent finding unexpected peaks between rows. The prior knowledge of the number of rows per microplot (*Row_number_prior*) is also used when identifying the peaks. The prior values of row and plant spacing are not always known precisely. Therefore, the row extraction pipeline (Figure 2) provides also updated and more accurate values of *Row_spacing_prior* for each session. Finally, each row is extracted using the fine-tuned value of the row width.

**Figure 1:**
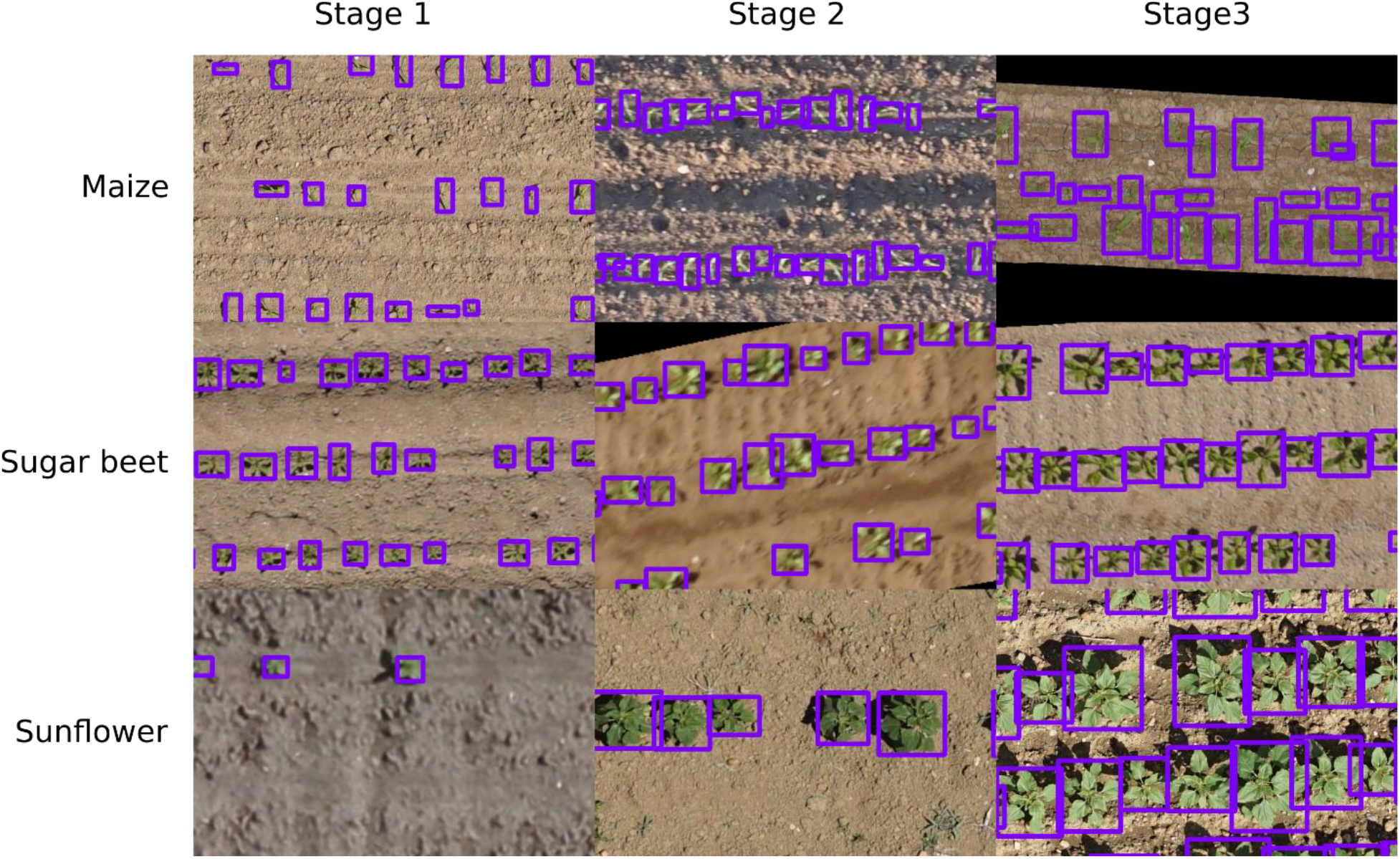
**Samples of images for the three-development stage. All images were resampled to 0.25mm.px**^**-1**^. **The bounding boxes were drawn interactively around the plants**.

**Figure 2.**
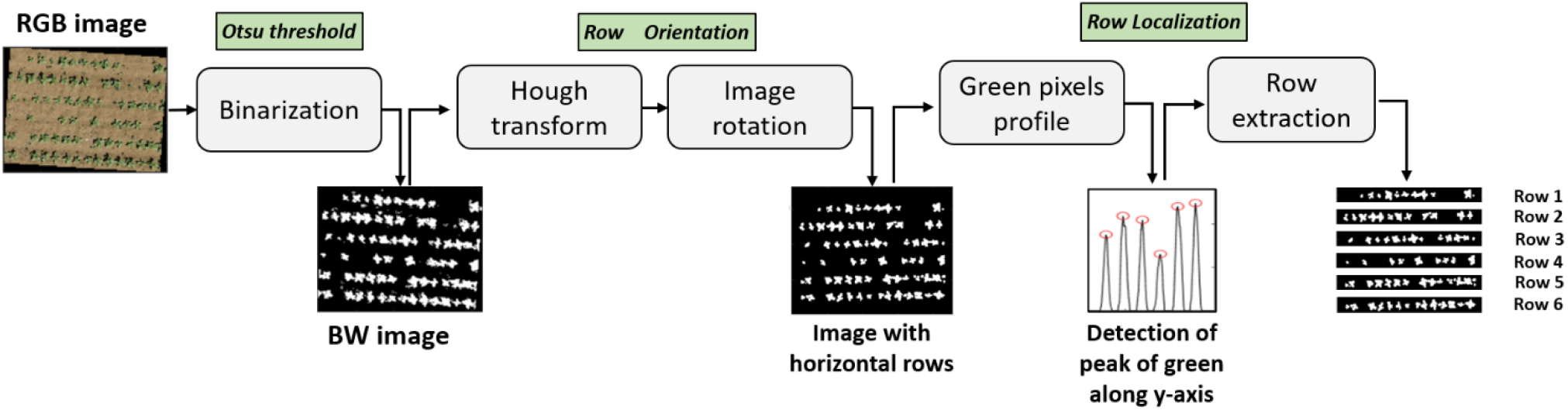
**Flowchart of the rows extraction process from the original RGB image**.

##### 2.2.1.2 Plant identification with an object-based method

After the row extraction, the algorithm individualizes the objects (groups of connected pixels) in the image and classifies them as plants or weeds. Weeds are eliminated based on the distance to the row center. If the centroid of an object is located at a distance larger than a threshold value (*Minimum_distance_to_row)*, it is considered as a weed. The threshold value is expressed in relative value to the row spacing and set to 0.25 (Table S2). Objects with dimensions along the row direction larger than the *Plant_spacing_prior* value (Table 2) are expected to include several plants. The number of plants contained in these big objects is derived from the number of peaks observed when summing the green pixels along the row direction, where a peak may correspond to a plant position. Further, the number of plants found by the number of peaks is crosschecked with the expected number of plants computed by dividing the extension of the object by the *Plant_spacing_prior* value. Results are illustrated in Figure 3 for the two objects on the right of the bottom row.

**Figure 3:**
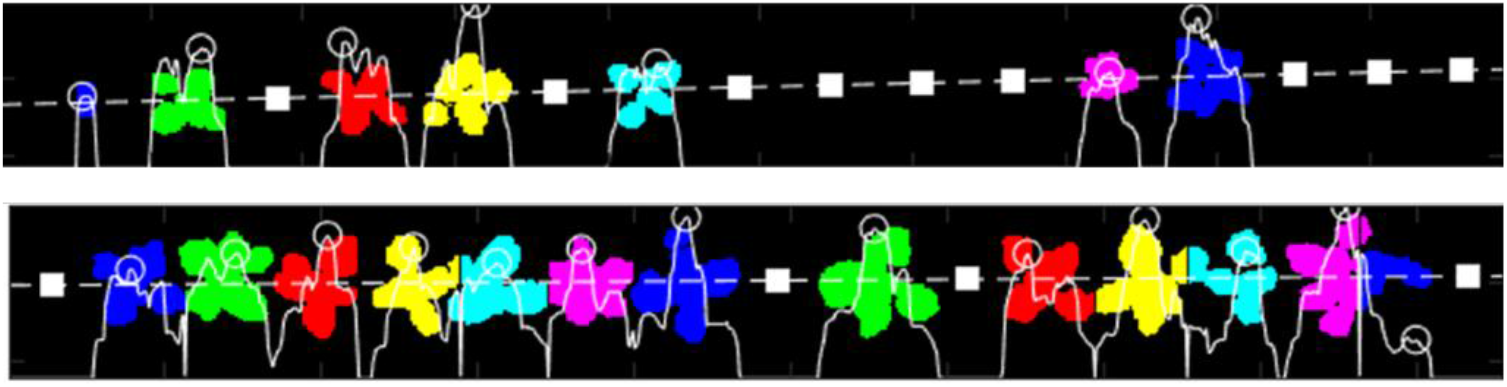
**Typical output of the HC algorithm illustrated for two sugar beet rows. The dashed white line indicates the row. The white curve represents the profile of number of green pixels perpendicular to the row, with peaks identified by a circle. The object-based method is illustrated by the colors assigned to each identified plant. Note that big objects have been split into individual plants (bottom row, the four last plants) and isolated plant parts have been reconnected to form a single plant (top row, fourth plant starting from the left). The white squares correspond to the position of missing plants**

Finally, some objects may be located too close together to be considered as separate plants because these objects correspond to several parts of the same plant. Figure 3 illustrates it with the second plant starting from the left on the top row, where a leaf and the main plant are separated. If the distance between the centroids of the closest object is smaller than the maximum acceptable distance, *Big_plants_tolerance x Plant_spacing_prior*, the two objects are merged as a single plant. Table S2 in the supplementary materials presents the value used for each parameter. The centroid (center of mass of the object), and the bounding box (smallest rectangle that contains all object’s pixels) of the objects are finally computed.

#### 2.2.2 Deep-learning method

##### 2.2.2.1 Model architecture

An object detection method was selected to predict the bounding box around each plant. This information can then be used to derive more traits to characterize every individual plant. Object detection is a fast-growing area within DL techniques since the emergence of networks such as R-CNN (Regions with Convolutional Neural Network, (Girshick et al., 2013)) or SSD (Single Shot Detector, (W. Liu et al., 2016)). Most DL object detection models fall into one-stage or two-stage models. In the one-stage model, the object is localized and categorized in a single step. In the two-stage model, a first stage detects possible objects, and a second stage categorizes them. The Faster-RCNN two-stage model (Ren et al., 2015) is used because it performs well in the context of plant phenotyping. Madec et al. (Madec et al., 2019) used it successfully for counting wheat heads. It allows also to analyze the nature of the possible errors by visualizing them.

Faster-RCNN can be implemented in many forms which can influence the final results. We use the implementation made by the mmdetection library (Chen et al., 2019). It contains many detectors, and is written upon PyTorch (Paszke et al., 2019). The default implementation of the library is used and contains a Feature Pyramidal Network (FPN) (Lin et al., 2017), which differs from the original paper (Ren et al., 2015). It is used to provide object proposition at different scales. A ResNet-34 model (He et al., 2015) was used as the backbone network because it offers a good compromise between accuracy and speed of training. The backbone extracts the deep features which are used by the Region Proposal Network (RPN) to detect potential objects which are then classified as crop or background. All other architectural details are given in the code (https://github.com/EtienneDavid/plants-counting-detection). We also choose to train one model by crop as preliminary tests show lower performances when mixing the three crops.

##### 2.2.2.2 Pre-processing and data augmentation

The input image size of the network is set to 512 × 512 pixels to match memory constraints during training. However, images from the microplots are larger. A preprocessing step first splits them randomly into patches of 512 × 512 pixels. For each session in the training dataset, 100 patches were randomly selected which results in a total of 900 patches to train the model for each crop over the nine available sessions. Randomly sampled patches provide more diversity than evenly sampled ones. During the training process, data augmentation is applied to extend the diversity of images. The complete data augmentation pipeline is a set of geometric distortions (Random rotation, Random Translation, Random Shear), blur (Gaussian Blur), noise (Gaussian noise) and colorimetric augmentation (Random hue value, Random contrast). At each iteration, a set of transformation is randomly drawn with random parameters so each batch is unique. The range of possible parameters were chosen so that the resulting image still look realistic. All data augmentation details are given in the code. Once trained, the model is applied to all the patches. Predictions from the overlapping patches are finally merged together by using the Non-Max-Suppression algorithm (Ghosal et al., 2019) with an Intersection over Union (IoU) threshold of 0.70.

#### 2.2.3 Hybrid method

DL methods detect individual plants based on many features automatically extracted while HC methods exploit expert prior knowledge on the sowing pattern to eliminate plants located at a non-expected position between rows. We propose therefore a hybrid method that combines the benefits of both HC and DL ones. The DL method is first applied to detect plants. Then, the HC method presented earlier is used to identify the row position and eliminate all remaining weeds corresponding to plants with centroids located at a larger distance to the row than a threshold value *distance_to_row* (Table S2).

### 2.3 Evaluation strategy for plant detection

#### 2.3.1 Strategies for training and evaluation

Detection models were developed and evaluated independently for each crop. DL method requires an extensive training dataset that should represent the expected diversity of situations. Due to the limited number of labelled images, two strategies are defined: “Out-Domain” and “In-Domain”. “Out-Domain” is the more rigorous strategy where the performances of the DL method are evaluated over sessions not used during the training process. For each crop and each stage, two sessions were used for training and the remaining one for testing. This allows to balance the stages between the training and testing datasets. A three-fold cross-validation strategy that exploits all sessions while providing relatively independent test cases is used. Three different models were trained for each crop using six sessions, representing about 3800 plants, and tested on the remaining three sessions representing around 1900 plants. The “In-Domain” strategy is based on adding few images randomly selected in the testing datasets to the training dataset. It aims at reducing possible lack of representativeness in the training dataset. The same three-fold cross-validation process was used for each crop, except that 1/3 of the 600 plants used previously as testing datasets were added to the training dataset. The remaining 2/3 images (400 plants) are used to evaluate the performances of the models for each crop. The same test dataset (1200 plants corresponding to the 400 test plants for each of the three test sessions) is finally used to compare the Out-domain and In-domain approaches. The approach is summarized in Figure 4.

**Figure 4:**
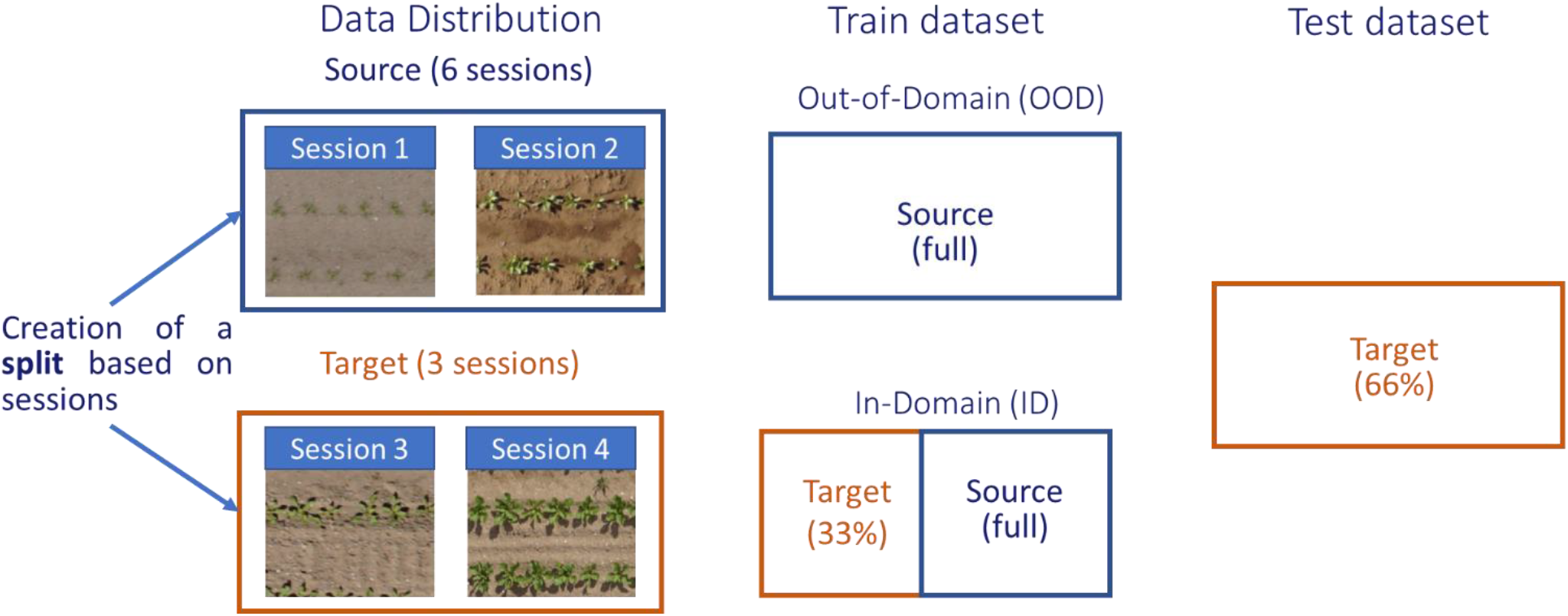
**Presentation of the strategy for training and evaluation. For each fold, we select 6 sessions as the training distribution and 3 as the target distribution. The test datset is made of 66% of the target distribution**.

#### 2.3.2 Evaluation metrics

##### Detection

The “Centroid matching strategy” (C_MS) is used to evaluate whether a plant was correctly detected. The C_MS is based on the distance between the centroids of the plants. If the distance between centroids of a detected plant and the closest labelled one is smaller than *Plant_distance_prior* / 2 it is considered as true positive (TP). Otherwise, it is a false positive (FP). If a labelled plant has no detected plant within a distance smaller than *Plant_distance_prior* / 2, it is a false negative (FN). TP, FP and FN are used to construct the confusion matrix (Equation 2).

**Equation 2: Presentation of the confusion matrix. Please note that in detection, there is no True Negative (TN)**

| Total population = Number of Ground Truth positive |  | Prediction |  |
| --- | --- | --- | --- |
|  |  | Predicted Positive (Box) | Predicted Negative (No box) |
| Ground truth | Positive (Box) | True Positive (TP) | False Negative (FN) |
|  | Negative (No box) | False Positive (FP) | True Negative (TN) |

The plant detection performance was quantified per session with the terms of the confusion matrix normalized by the number of labelled plants (TP+FN) for easier comparison between crops and stages, which correspond to rates of TP (TPR), FP (FPR) and FN (FNR). The accuracy is also used, defined as TP/(TP+FN+FP). DL method produces a confidence score for each predicted BB. A box is considered as a prediction for the DL and HY methods if its score is above 0.5.

##### Plant density

Plant density (PD) was calculated by dividing the number of plants in the microplot by its area. The area is computed as the number of rows multiplied by the row spacing and the row length. The relative root mean square error (rRMSE) is used to compare the estimated and the reference PD values and assess the accuracy of the method. The accuracy levels were split into four classes to better assess the robustness of the method. A rRMSE<5% was considered as good, between 5%<rRMSE<10% as satisfactory, between 10%<rRMSE< 20% as poor, and rRMSE>20% as very poor. The percentile of microplots belonging to each class was therefore used to evaluate the robustness of the methods.

**Equation 3: Definition of the rRMSE for one session of acquisition**

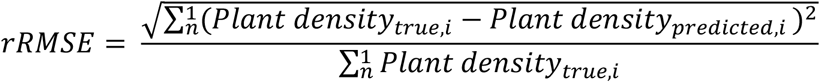

##### Influence of conditions

Tests were further conducted to evaluate the impact of the four qualitative factors (crop type, development stages, weeds, and soil type) and the impact of the four quantitative factors (sowing density, plant size, original resolution, and blurriness). For the qualitative factors, an ANOVA study is conducted, and for the quantitative factors a Pearson test is conducted. Both modalities were implemented with the python statsmodel library. For both tests, the p-value is calculated to evaluate the impact of the agronomical conditions on the final results.

## 3 Results

### 3.1 Detection

Detection performances are very different depending on the crops (Table 4 and Figure 5). Detection of maize plants appears difficult for the three methods and particularly for HC with a low TPR and a high FNR (Table 4). However, a high FNR is also observed for the first development stage with the HC method. A large variability between the three instances of the three-fold cross validation is observed for this early stage (Figure 5), probably due to the variability in image quality. Marginal differences are observed between DL and HY methods. They both show relatively balanced FPR and FNR. This results into accuracy values between 0.77 to 0.80 with little variation between stages (Table 4). However, a larger variability across the three instances of the three-fold cross validation is observed for the late stage (Figure 5)

**Table 4:** **Terms of the confusion matrix for the three methods the three crops, and the three stages. True Positive Rate (TPR), False Positive Rate (FPR), and False Negative Rate (FNR) are displayed. N is the true number of plants (N=TP+FN). Green color corresponds to good metrics values (high for TPR, low for FPR and FNR), and red for poor metrics values (low for TPR, high for FPR and FNR)**.

| Crop | Stages | N | TPR |  |  | FPR |  |  | FNR |  |  | Accuracy |  |  |
| --- | --- | --- | --- | --- | --- | --- | --- | --- | --- | --- | --- | --- | --- | --- |
|  |  |  | HC | DL | HY | HC | DL | HY | HC | DL | HY | HC | DL | HY |
| Maize | 1 | 1669 | 0.61 | 0.88 | 0.86 | 0.27 | 0.12 | 0.07 | 0.39 | 0.12 | 0.14 | 0.56 | 0.79 | 0.80 |
|  | 2 | 1658 | 0.70 | 0.92 | 0.92 | 0.03 | 0.18 | 0.16 | 0.30 | 0.08 | 0.08 | 0.68 | 0.78 | 0.79 |
|  | 3 | 1930 | 0.70 | 0.88 | 0.86 | 0.05 | 0.15 | 0.14 | 0.30 | 0.12 | 0.14 | 0.68 | 0.77 | 0.77 |
| Sugar beet | 1 | 1825 | 0.95 | 0.98 | 0.98 | 0.04 | 0.01 | 0.01 | 0.05 | 0.02 | 0.02 | 0.93 | 0.97 | 0.97 |
|  | 2 | 1948 | 0.95 | 0.99 | 0.99 | 0.01 | 0.03 | 0.03 | 0.05 | 0.01 | 0.01 | 0.93 | 0.97 | 0.97 |
|  | 3 | 1874 | 0.94 | 0.99 | 0.99 | 0.06 | 0.04 | 0.04 | 0.06 | 0.01 | 0.01 | 0.88 | 0.95 | 0.95 |
| Sunflower | 1 | 1603 | 0.80 | 0.87 | 0.86 | 0.17 | 0.06 | 0.04 | 0.20 | 0.13 | 0.14 | 0.75 | 0.80 | 0.81 |
|  | 2 | 1856 | 0.82 | 0.94 | 0.94 | 0.15 | 0.08 | 0.07 | 0.18 | 0.06 | 0.06 | 0.72 | 0.87 | 0.88 |
|  | 3 | 1759 | 0.86 | 0.97 | 0.97 | 0.42 | 0.43 | 0.21 | 0.14 | 0.03 | 0.03 | 0.61 | 0.74 | 0.81 |

**Figure 5:**
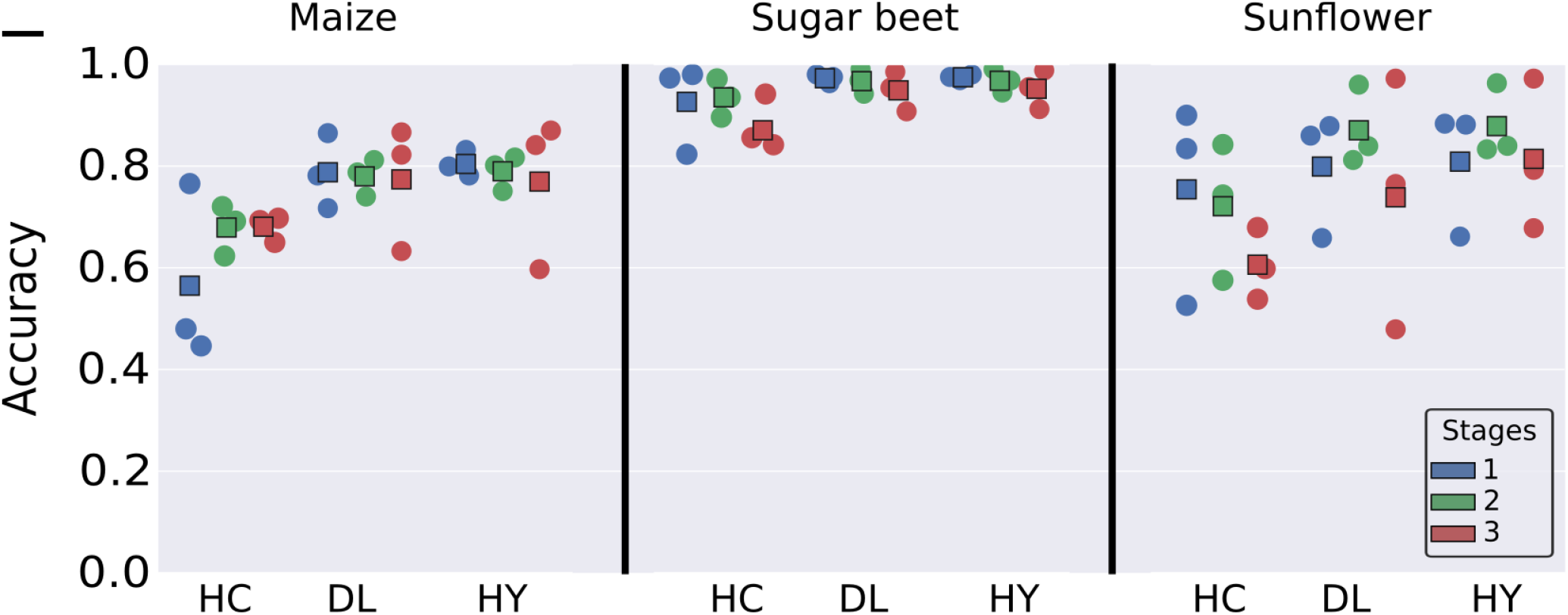
**Accuracy for all methods and crops. For each crop and method, the stages are represented by a specific color. Each point corresponds to a test session used in the three-fold validation process. The squares represent the average of the three points**.

### 3.2 Counting

The HC method provides the poorest performances for maize plant density estimation, with rRMSE generally higher than 0.2 (Figure 6), which is consistent with the poorer detection performances (Figure 5). Image acquisition during the early stages tends to degrade the performances conversely to what was observed for the detection (Figure 5). This may be explained by the unbalance between false positives and negatives observed for the early stages (Table 4). Marginal differences are observed between DL and HY methods for maize where weeds were not the main issue.

**Figure 6:**
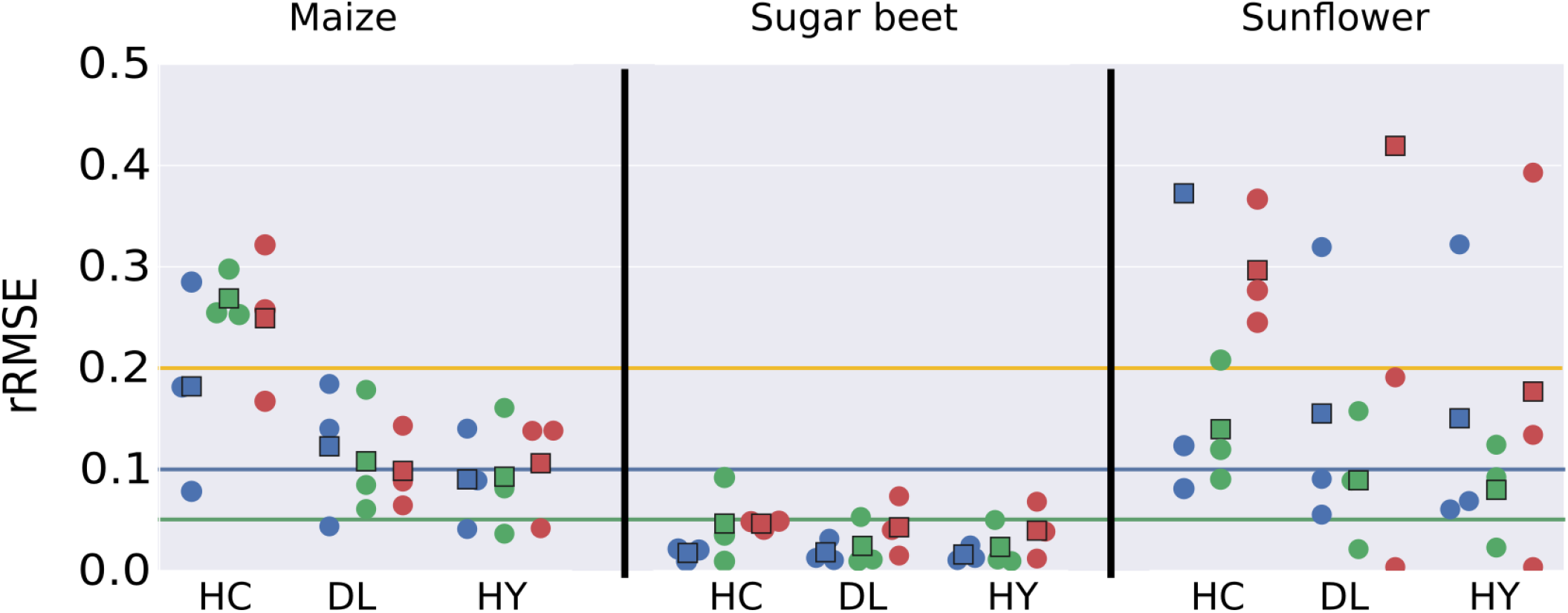
**rRMSE for plant density estimation for all methods crops, and stages. Results obtained over the testing dataset. For each crop, method and stage, the three instances (corresponding to three testing sessions) of the three-fold cross validation process are displayed as colored dio sks, while the corresponding average is represented by a colored square. Colors correspond to stages. The rRMSE threshold values to acceptable level of performance (green: very good, blue: good, orange: acceptable)**

### 3.3 Out-Domain against In-Domain results

The “Out-domain” strategy used previously was compared here to the “In-domain” one where 1/3 of the images of the initial testing sessions were used to finetune the model. Performances are evaluated on the remaining 2/3 images of the initial testing sessions to keep some independence between the training and test datasets. Results show that the additional images used in the training process and having similar characteristics as those in the testing dataset decreased significantly the rRMSE for all crops (Figure 7). Training with the In-domain strategy reduces the variability of performances across sessions. The 5% rRMSE value is reached for all crops except maize, where performances are anyway close to this target. Plant overlapping and the small leaf size makes the DL method for maize more challenging. However, there are still some outliers for Maize and Sunflower, corresponding to Pleinefougeres_2019_1 and Epoisses_2019_1 sessions. The images of these two sessions are highly blurred (Table 3) explaining most of their poor detection performances. A large part of this performance can be attributed to the elimination of almost all weeds by the DL methods, without the need of the HY correction, which have learned the pattern of the weeds, instead of relying on the location, and a better recognition of the plants.

**Figure 7:**
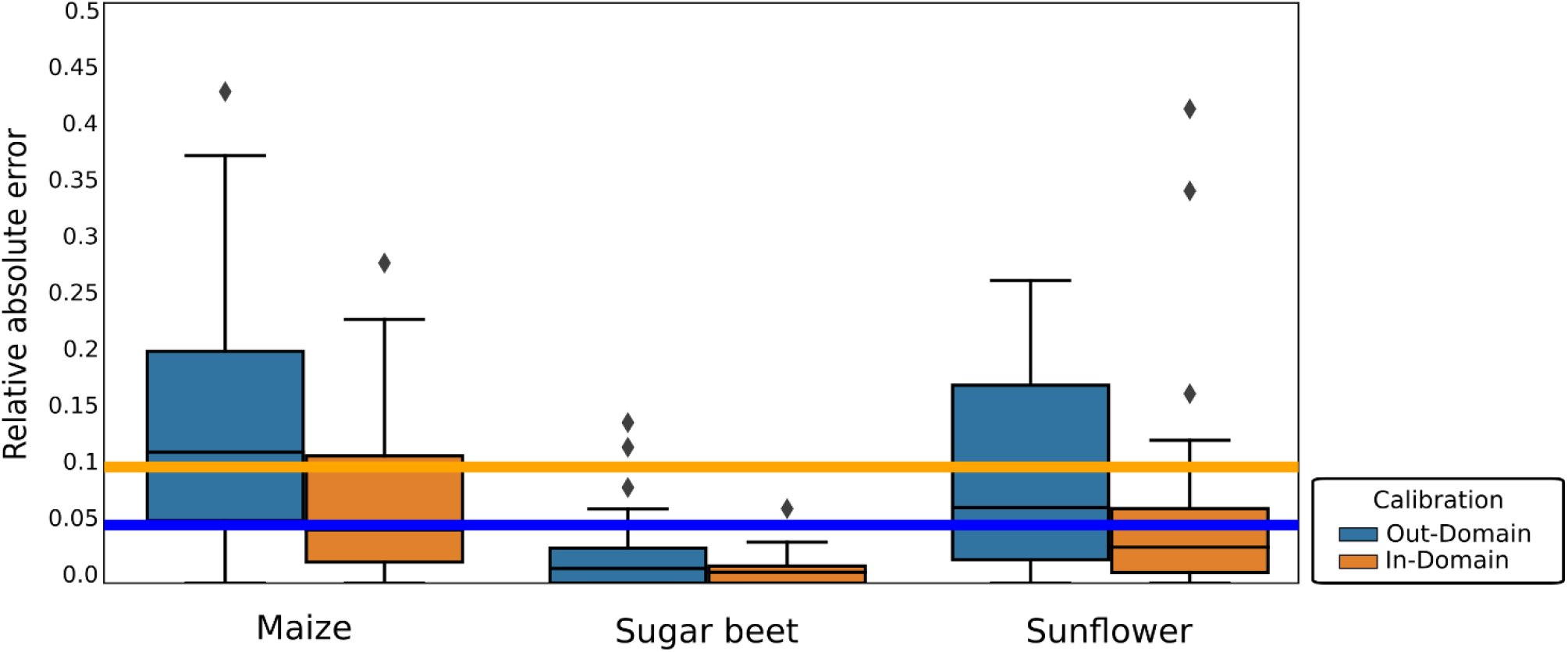
**Distribution of relative absolute error for each microplots for the Out-Domain and In-Domain approaches for DL. Box-plot representation where the black horizontal bar represents the median, the box represents ±25%, the whiskers while the whiskers extend to the the lowest (highest) data point still within 1.5 interquantile range of the lower (upper) quartile. Diamonds are outliers. 1 outlier for Out-domain Maize and 3 outliers for Out-Domain Sunflower are above 0.5 and are not presented on the graph**.

## 4 Discussion

### 4.1 DL and HY methods detect better plants than the HC one

Several factor can explain the variability of the results: the small size of the plants that overlap, resulting into groups of overlapping plants that are interpreted as a single plant (Figure 5b), or to poor threshold values determined by the Otsu method for the green segmentation used in the first step to identity objects (Figure 5a), due to the poor quality of the green segmentation where background artifacts such as small rocks or crop residues were interpreted as plants (Figure 5g). Also, in some case a high FPR is mostly explained by possible confusion between plants and their shadows or soil artifacts (Figure 5c) while FNR is explained by the small size of the plants that are difficult to detect (Figure 5d).

Detection of sugar beet plants appears to be much easier, with performances similar between the three methods. The sugar beet crops better verify the assumptions described in 3.2.1. The plots were not infested by weeds (Table 4), which seems to be an important explanation for the success of all methods. A small FPR is observed for the three methods, particularly for the latest stage, which explains the decrease in accuracy (Table 4). This is due to difficulties when plants are overlapping (Figure 5e). Slightly higher FNR is observed for HC corresponding to non-detected plants in the case of small plants and image of poor quality. This is also observed with DL for the very early stages (Figure 5f). The variability across the three instances of the three-fold cross validation is also small (Figure 4). Marginal differences are observed between DL and HY methods mostly because of the good control of weeds.

Detection of sunflower plants shows accuracy values intermediate between maize and sugar beet (Table 4 and Figure 4). The HC shows lower TPR and higher FPR and FNR as compared to DL and HY. In the late stage, the HC shows very high FPR corresponding to problems of plant separation when they are overlapping. Further, the weeds close to the row line are not well eliminated and confounded with plants (Figure 5g). Similar problems are observed for the DL method, with weeds confounded with the crop. However, the HY methods allows to eliminate part of the weeds that are located in between rows (Figure 5h). and HY shows high and similar TPR (Table 4). However, a high FPR is also observed for the first stage with the HC method, due to the poor quality of the green segmentation where background artifacts, such as small rocks or crop residues, were interpreted as plants (Figure 4g). Conversely, high FPR are observed for the late stage where DL shows difficulty to detect plants in a group of overlapping ones and confounds weeds with the crop. A large variability between the three instances of the three-fold cross validation is observed for sunflower (Figure 4). It is explained by a high degree of heterogenety in the microplots and between them, as well as between sessions.

**Figure 8:**
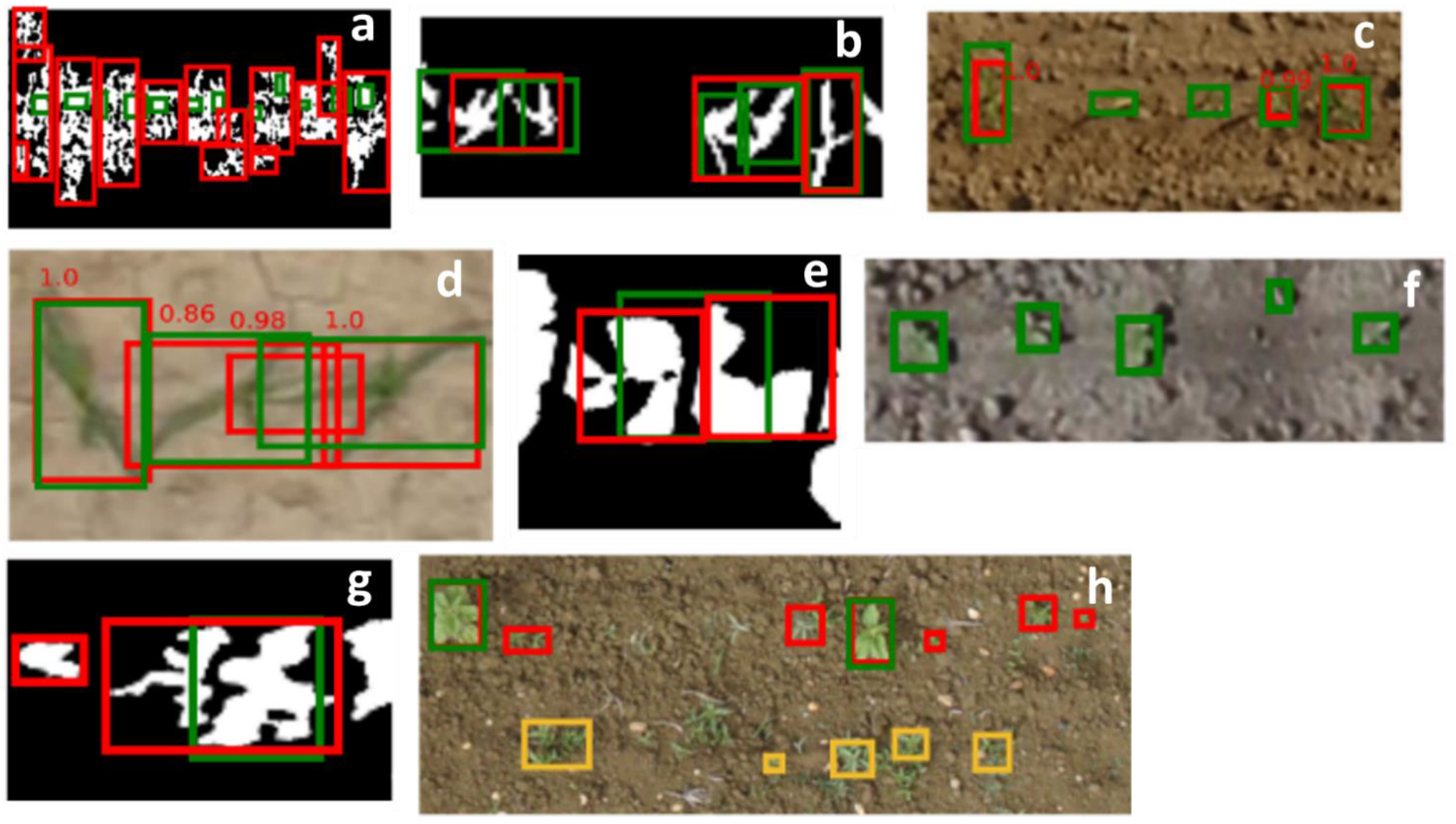
**Possible detection errors for HC and DL methods. The green BBs correspond to the labelled plants. The red BBs correspond to the detected plants and yellow boxes correspond to weeds detected as crop. RGB images are displayed for the DL method. BW images are displayed for the HC method. a, b, c, d corresponds to maize, e, f, to sugar beet and g, h to sunflower**.

Image quality appears therefore mandatory for HC methods to get a good segmentation. The HC methods appears also limited to eliminate weeds on the rows and to separate efficiently the overlapping plants. DL methods are similarly limited in separating crops from weeds, with confusions made mostly on unseen type of weeds (Figure 5h). However, the HY methods allows to eliminate part of the weeds. The DL methods also show some difficulties in detecting plants when they are small or when their shadows or other soil artifacts such as cracks are present. Nevertheless, our DL methods seems to outperform the HC ones in most cases.

Tests were further conducted to evaluate the impact of the four qualitative factors (crop type, development stages, weeds, and soil type) using the p-value computed from a variance analysis. Results show (Table 5) that crop-type is an important factor (p_value smaller than 0.05) for HC and HY, while weeds are important for HC and DL, and soil-type for HC. However, the low number of examples (27 sessions in total), and the non-evenly distribution of the several factors (for instance most examples of high levels of weed infestation are found in sunflower sessions only) prevents from drawing final conclusions. The impact of the four quantitative factors (sowing density, plant size, original resolution, and blurriness) were also evaluated using a Pearson test. It reveals (Table 5) that no factors appear significant (p-value smaller than 0.05), while the lowest p-values are observed for the sowing density and plant size that are closely related to the crop type.

**Table 5:** p-values computed from an ANOVA for the qualitative factors and Pearson test for the quantitative factors.

| Factors | Type | HC | DL | HY |
| --- | --- | --- | --- | --- |
| Crop type | qualitative | 0.009130** | 0.127550 | 0.032050** |
| Development stage | qualitative | 0.857810 | 0.479530 | 0.643620 |
| Weed infestation | qualitative | 0.032610** | 0.001600** | 0.074540 |
| Soil type | qualitative | 0.026430** | 0.781090 | 0.830650 |
| Sowing density | quantitative | 0.067379 | 0.076542 | 0.091679 |
| Original resolution | quantitative | 0.905626 | 0.572383 | 0.616534 |
| Plant size | quantitative | 0.791437 | 0.064765 | 0.211019 |
| Blurriness | quantitative | 0.111743 | 0.562775 | 0.501980 |

### 4.2 Plant density is better estimated with DL and HY methods

All the methods reach good performances (rRMSE<0.05) for sugar beet, with even better performances for the two first stages when plants are easily identified and weeds not too developed (Figure 6). The poorer detection performances noticed earlier for HC (Figure 4) do not impact the density estimation because the FPR is well compensated by the FNR.

Sunflower shows more variability between sessions and stages, with rRMSE around 0.1 for the intermediate development stage showing better performances than the early one and moreover than the late one (Figure 6). The models for sunflower are very poor for the session 3_auzeville_2019_5 (Figure 6), mainly because of weed infestation. DL performs better than HC while HY improves marginally the performances for the two early stages, but significantly for the late stage where significant weed infestation was observed.

Overall, our results show lower performances than those of the studies where the training and testing datasets were not independent. For maize detection accuracy between 0.93 and 0.96, and relative counting error around 1.5%, were reported (Quan et al., 2019; Varela et al., 2018) while none of our methods achieve such performances. Similar range of results are obtained on rapeseed (counting error of 6.83%) (Zhao et al., 2018), or safflower with rRMSE approximately under 5% (Koh et al., 2019). However, our results with DL and HY are comparable to studies keeping the training and test datasets independent; on maize Gnädinger and Schmidthalter (Gnädinger and Schmidhalter, 2017) reports a counting error of +/- 15%. The HC approach applied when its main assumptions are verified performs well and comparably to DL.

### 4.3 Adding few images from the test domain improves drastically the DL performances

The performances of DL methods are closely related to the number of images used in the training dataset and their representativity of the possible situations (Geirhos et al., 2020). DL method works very well for sugarbeet where all the images were relatively similar across sessions for each development stage. However, the acquisition conditions were quite different from the ones experienced in the other sessions for the sunflower on Epoisses_2019_1, explaining why the DL models had more difficulties to detect plants for this session. Note first that the plant density estimation performances (Figure 7) evaluated on a limited test data set (1200 images) are very consistent with the ones presented previously over the full test dataset including 1800 images (Figure 6). Overall, the addition of in-domain data largely outperforms the marginal gain observed with the HY method on few sessions.

Our results demonstrate that active learning techniques (Ghosal et al., 2019) could greatly improve DL model performances for these new sessions. A small sample of images coming from the new sessions to be processed have to be labelled to complement the training dataset, but more than quantity, it is uniquely due to the diversity: only 40m² of maize or sugarbeet, and between 50 and 100m² of sunflower have been added to the training dataset, leading to a dramatic increase of the performances which cannot be attributed only to the dataset size increase. These results demonstrate the importance of having a proper design of DL training dataset when proposing a new trait to get robust estimates as required by agronomists, breeders, and farmers.

Our results are consistent with those of previous studies: detection and density estimation performances are generally lower when the training and the test datasets are independent, i.e not coming from the same measurement sessions. Fernandez-Gallo (Fernandez-Gallego et al., 2020)report a rRMSE below 5%, Madec et al. (Madec et al., 2019) report a rRMSE of 15% on an independent test set. Similar drop in performances seems to happen in maize when comparing the results of Varela et al. (counting error of 1.5%) to those of Gnädinger and Schmidhalter (counting error of +/- 15%). The generalization potential of DL methods is high, requiring including more diverse situations in the training dataset at the expense of the tedious and expensive interactive labelling process. However, alternative techniques could be used to bypass this limitation, including data sharing between several organizations as this was done for the head counting problem (David et al., 2020). Data augmentation (Kuznichov et al., 2019) could also improve greatly the generalization performances of DL methods. It would consist in manipulating the quality of the images, while creating synthetic images where a wide diversity of plants and weeds would be placed over different backgrounds with variation in the development stages and sowing pattern.

## 5 Conclusion

This study was based on a comprehensive dataset covering three main crops, several growth stages and acquisition conditions. It will be open to the community on Zenodo (https://zenodo.org/record/4890370) to be possibly used as a benchmark for plant counting and detection from RGB images acquired from UAVs. Our results show that when the main assumptions on the sowing patterns are verified, simple HC methods can reach good enough performances to be used for applications as it was observed here for sugar beet. However, simple Deep Learning methods generally outperform the simple HC ones. Nevertheless, due to the large heterogeneity in terms of background, plant shape and phenological stages encountered across the wide collection of images considered, we demonstrated that the performances of the DL methods largely depend on the training and test datasets used. When the training domains used for the DL method are fully independent from the testing ones, the overall performances are reduced due to the failure of the model in a number of test cases poorly represented in the training dataset. Conversely, when adding few examples of images representative of the test domain, the performances increase drastically to reach those reported in most studies where training and test domains are not differentiated. Important gain in robustness could therefore be reached by including in the training dataset few images coming from the inference domains. Alternatively, a better understanding of the factors of variability between domains could constitute the basis to generate efficient data augmentation techniques that may even include synthetic images. An extended version of the dataset is needed to conclude on the main factors of error on plant counting with UAV. The hybrid method proposed to better eliminate weeds could be replaced efficiently by including images of the canopy where weeds were artificially incrusted.

## 6 Acknowledgments

The work received support from ANRT for the CIFRE grant of Etienne David, co-funded by Arvalis. The study was partly supported by several projects, including ANR PHENOME. Many thanks to the dataset contributors:

- Arvalis (maize) : Menainville, Nerac, Villedieu, Thenay, Castetis
- Hiphen Plant (maize) : Blois, Selommes, Ermine, Pleinefougère
- Institut Technique de la Betterave (ITB) : All sugarbeet datasets
- Terres Inovia (sunflower) : Epoisses
- INRAe (sunflower): Auzeville, Rivière

## 8 Supplementary material (to put in an external file for submission)

**Table S1.**
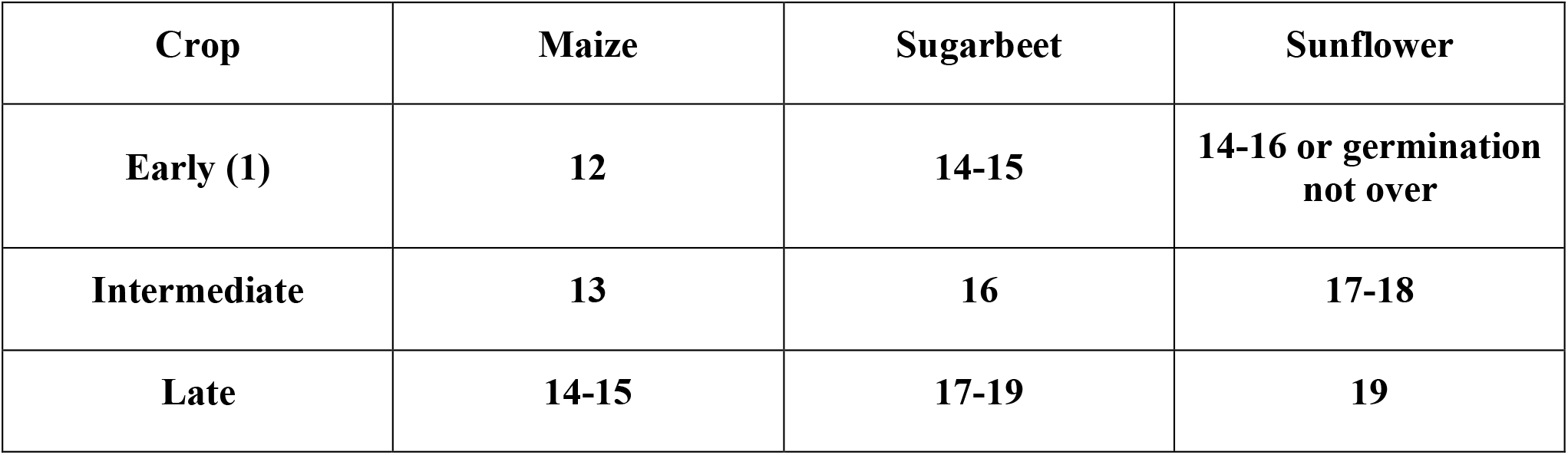
Correspondance between the “Early”, “Intermediate” and “Late stage” and the BBCH scale for each crop.

**Table S2.**
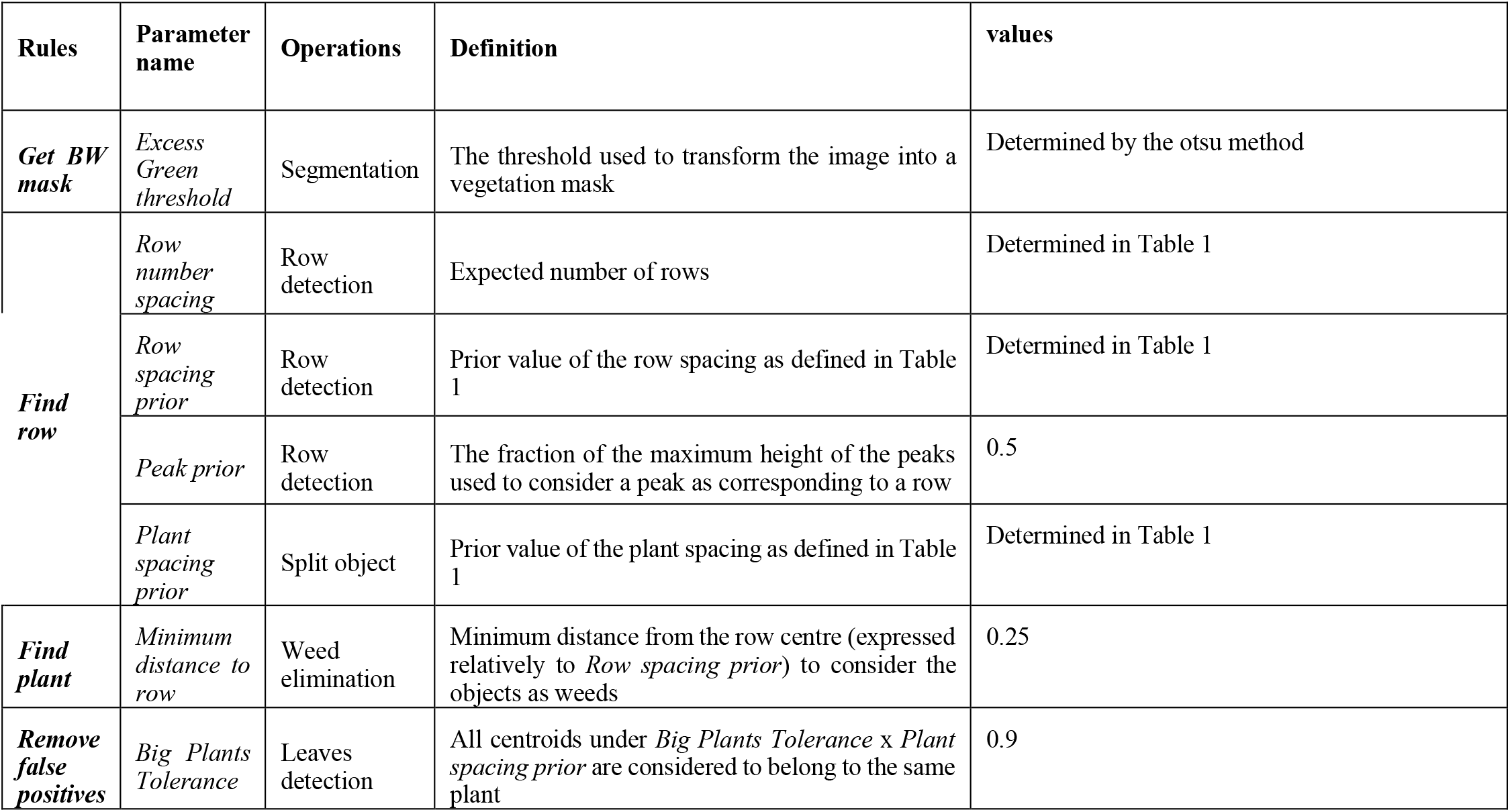
List of parameters used for row extraction and plant identification.

Figure S3: Justification of a centroid matching strategy Centroid matching strategy (C_MS) is preferred to the IoU one (IoU_MS)

The C_MS was initially compared with an intersection over union matching strategy (IoU_MS), which is more commin The IoU_MS is based on the Intersection over Union between the detected and labelled

BB with a standard threshold of 0.5. A detected plant is considered true positive (TP) if its IoU is larger than 0.5. Otherwise, it is a false positive (FP). If a labeled BB has no overlap with any detected BB, it is classified as false negative (FN).

The size of BB of plants detected by the HC method have different dimensions as compared to the labelled BB (Figure 4, left): The distribution of the size of BB for HC is gaussian, while that of labelled, DL and HY are very similar and skewed with significantly smaller BBs as well as larger ones. That means that the HC is missing small object with the IoU_MS. This resulted into lower values of accuracy computed with IoU_MS (Figure 4, right) because of a significant amount of mismatch between the predicted and reference BBs at IoU=0.5. Rather than adapting the IoU threshold level, the distance between centroids is preferred to evaluate the match between predicted and interactively labeled plants. The accuracy computed with C_MS (Figure 4, right) is significantly larger than that computed with IoU_MS, particularly for the low accuracy values as well as for the HC method for the reasons exposed above. Therefore, in the following, the centroid distance is used to compute the terms of the confusion matrix and the accuracy. Detailed metrics can be found in Table S2.

**Figure S3:**
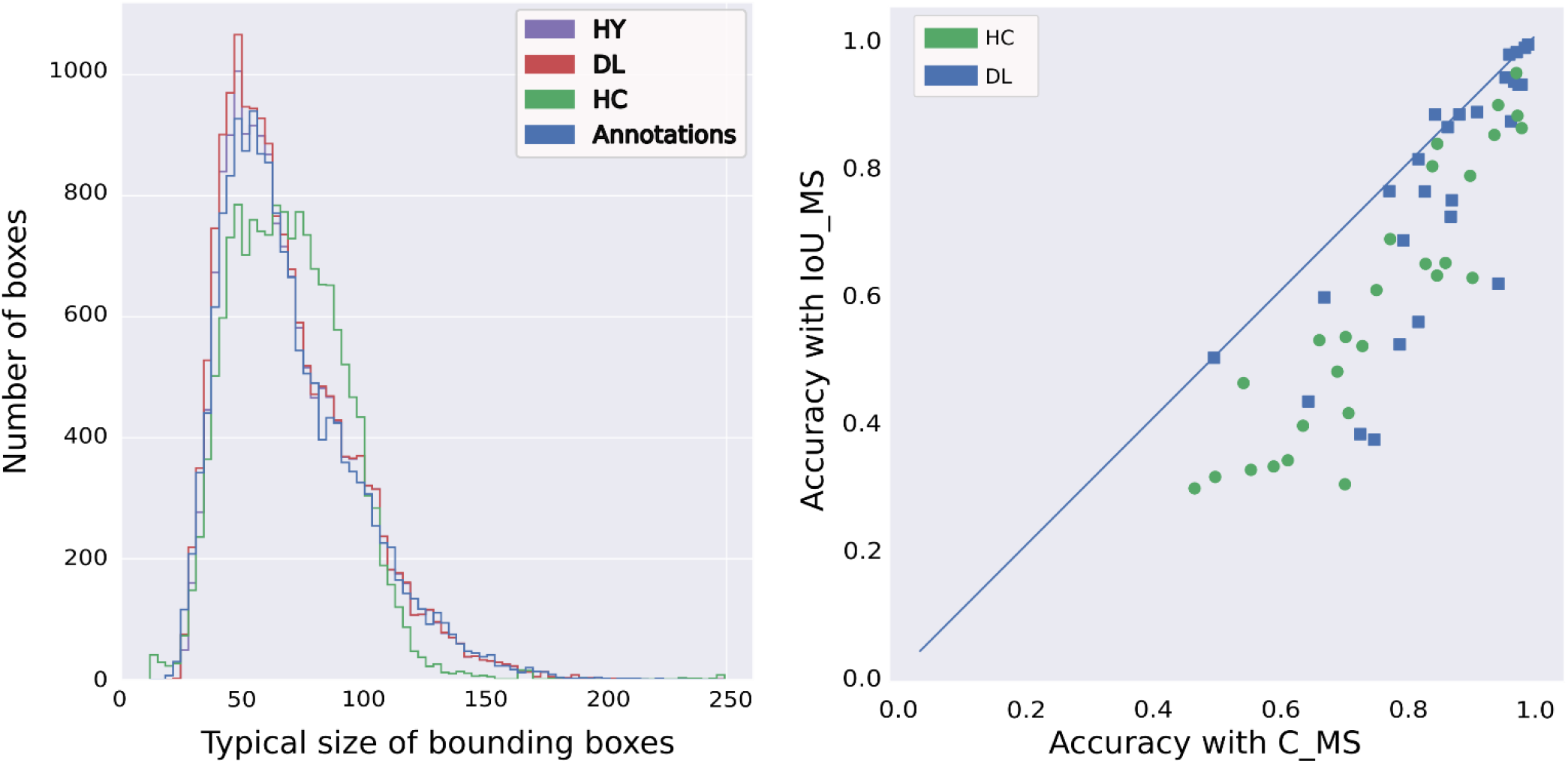
**Left: distribution of the typical size of BB annotated and those defined around the plants identified by the HC method. Right: comparison of Accuracy computed either with IoU_MS, and with C_MS for HC (green discs), and DL methods (blue squares)**.

**Table S4. Complete results for the three methods on all sessions. Accuracy, precision and recall are presented with the IoU matching strategy**.

## Notes

https://zenodo.org/record/4890370#.Yj3mgk2ZOUk

### Competing Interest Statement

The authors have declared no competing interest.

### Summary of Updates

This version is the results of many updates with reviewers

